# Biosynthetic proteins targeting the SARS-CoV-2 spike as anti-virals

**DOI:** 10.1101/2022.05.10.491295

**Authors:** Stéphanie Thébault, Nathalie Lejal, Alexis Dogliani, Amélie Donchet, Agathe Urvoas, Marie Valerio-Lepiniec, Muriel Lavie, Cécile Baronti, Franck Touret, Bruno da Costa, Clara Bourgon, Audrey Fraysse, Audrey Saint-Albin-Deliot, Jessica Morel, Bernard Klonjkowski, Xavier de Lamballerie, Jean Dubuisson, Alain Roussel, Philippe Minard, Sophie Le Poder, Nicolas Meunier, Bernard Delmas

## Abstract

The binding of the SARS-CoV-2 spike to angiotensin-converting enzyme 2 (ACE2) promotes virus entry into the cell. Targeting this interaction represents a promising strategy to generate antivirals. By screening a phage-display library of biosynthetic protein sequences build on a rigid alpha-helicoidal HEAT-like scaffold (named αReps), we selected candidates recognizing the spike receptor binding domain (RBD). Two of them (F9 and C2) bind the RBD with affinities in the nM range, displaying neutralisation activity *in vitro* and recognizing distinct sites, F9 overlapping the ACE2 binding motif. The F9-C2 fusion protein and a trivalent αRep form (C2-foldon) display 0.1 nM affinities and EC_50_ of 8-18 nM for neutralization of SARS-CoV-2. In hamsters, F9-C2 instillation in the nasal cavity before or during infections effectively reduced the replication of a SARS-CoV-2 strain harbouring the D614G mutation in the nasal epithelium. Furthermore, F9-C2 and/or C2-foldon effectively neutralized SARS-CoV-2 variants (including delta and omicron variants) with EC_50_ values ranging from 13 to 32 nM. With their high stability and their high potency against SARS-CoV-2 variants, αReps provide a promising tool for SARS-CoV-2 therapeutics to target the nasal cavity and mitigate virus dissemination in the proximal environment.

**Author Summary:** The entry of SARS-CoV-2 in permissive cells is mediated by the binding of its spike to angiotensin-converting enzyme 2 (ACE2) on the cell surface. To select ligands able to block this interaction, we screened a library of phages encoding artificial proteins (named αReps) for binding to its receptor binding domain (RBD). Two of them were able to bind the RBD with high affinity and block efficiently the virus entry in cultured cells. Assembled αReps through covalent or non-covalent linkages blocked virus entry at lower concentration than their precursors (with around 20-fold activity increase for a trimeric αRep). These αReps derivates neutralize efficiently SARS-CoV-2 β, γ, δ and Omicron virus variants. Instillation of an αRep dimer in the nasal cavity effectively reduced virus replication in the hamster model of SARS-CoV-2 and pathogenicity.

## Introduction

With up to 6 million deaths worldwide in less than two years, the COVID-19 crisis has demonstrated the necessity to better understand and fight the spread and transmission of respiratory viruses. Such knowledge will help to develop new efficient anti-viral strategies to mitigate future epidemics and pandemics. SARS-CoV-2 infection starts in the nasal cavity, the virus replicating at high titres in the olfactory epithelia before reaching the lower respiratory tract where it induces the main pathology [1]. Infection of the olfactory epithelium leads to massive damage which may explain the high prevalence of smell loss (anosmia) during the COVID-19 pandemic and to environmental dissemination to infect conspecifics [2],[3]. Blocking virus multiplication with antivirals delivered in the nose and the upper respiratory tract might therefore allow therapeutic benefit and prophylactic protection.

Series of human neutralizing monoclonal IgG antibodies and nanobodies/VHH fused to a Fc IgG domain able to inhibit SARS-CoV-2 infection have been produced and tested for systemic treatments, but their efficacy by delivery in the nose may not be optimal, due to a poor stability in the nasal cavity environment. Their firmness upon nebulization and aerosolization will be also a main issue for their use as therapeutics. Furthermore, their large-scale production should be economically not affordable in eucaryotic systems and technically difficult to achieve in prokaryotes [4].

As an alternative approach to VHH and antibodies, a family of artificial proteins, named αRep, was designed to provide a hypervariable surface on αRep variants [5]. αReps are thermostable proteins constituted by alpha-helicoidal HEAT-like repeats (31-amino acids long) commonly found in eukaryotes [6] and prokaryotes [7], including thermophiles. Sequences of homologs form a sharply contrasted sequence profile in which most positions are occupied by conserved amino acids whereas other positions appear highly variable generating a versatile binding surface (**Fig. 1**). A large αRep library has been assembled and was demonstrated on a wide range of unrelated protein targets to be a generic source of tight and specific binders. Thus, αReps were previously selected as interactors of HIV-1 nucleocapsid and to negatively interfere with virus maturation [8].

**Fig 1.**
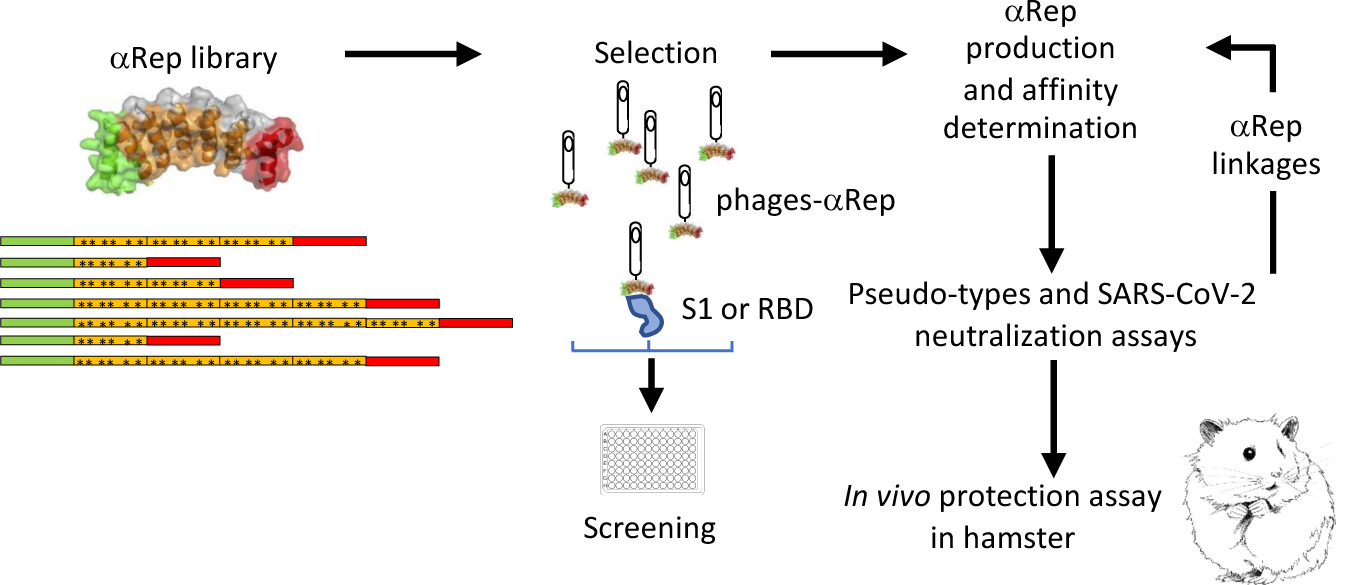
Selection and characterization of anti-spike αReps. Screening an αReps phage library allowed the identification of several binders specific of the RBD of the SARS-CoV-2 spike protein. Their binding affinity for the S1 domain was measured by biolayer interferometry. The neutralization activity of selected αReps was evaluated using a pseudo-typed S SARS-CoV-2 neutralization assay and a SARS-CoV-2 infection assay. Competitive binding assays were carried out by BLI to identify αReps recognizing non-overlapping binding sites. Then, αRep derived constructs followed the same characterization steps than their single counterparts. The protective potency of the best candidate was analyzed *in vivo* in the golden Syrian hamster model.

As for all coronaviruses, the SARS-CoV-2 spike (S) protein mediates virus entry to permissive cells. The S protein is a trimeric class 1 fusion protein that binds to its cell receptor, angiotensin converting enzyme 2 (ACE2), before undergoing a dramatic structural rearrangement to fuse the host-cell membrane with the viral membrane [9], [10]. Fusion is triggered when the S1 subunit binds to a host-cell receptor via its receptor binding domain (RBD). In order to bind to the receptor, the RBD undergoes articulated movements that transiently expose or hide its surface associated to the binding to ACE2 [11]. The two states are referred to as the “down” and the “up” conformations, in which down corresponds to a state incompetent to receptor binding and up to a state allowing receptor recognition. Due to its key function in the virus cycle, the RBD represents a target to identify binders that block interaction with the host-cell receptor or movements of the RBD between the down to up conformations [12].

Most SARS-CoV-2 infected people presents serum neutralizing antibody activity against the RBD indicating its immunodominance [13]. To reduce antibody binding, the viral evolution has led to the appearance of specific escape mutations in the RBD making current antibody-based treatments rapidly less effective [14].

Here, we first obtained a series of αReps specific of the receptor binding domain of the spike of SARS-CoV-2. These ligands display high affinities and blocked SARS-CoV-2 infection *in vitro*. The assembly of αRep through covalent and non-covalent linkages lowers the neutralisation EC_50_ to the 10 nM range. The αRep F9-C2 fusion protein instilled in the nose was found to limit virus replication and inflammatory response in a hamster model of SARS-CoV-2 infection. Furthermore, the F9-C2 fusion protein and a C2 homotrimer were found as potent inhibitors of SARS-CoV-2 variants including the antigenically distant omicron variant.

## Results

### Selection of αReps binders of the SARS-CoV-2 receptor binding domain

An overview of the selection process to generate anti-SARS-CoV-2 αReps specific of the spike is shown in **Fig. 1**. In order to select binders blocking SARS-CoV-2 entry into cells, the RBD (amino acids 330 to 550 of the spike S sequence) was used as a bait for screening. The phage display procedure included three rounds of panning followed by a screening step by phage-ELISA on individual clones. Nucleotide sequencing allowed the identification of >20 independent clones that were retained for further analyses (selected αRep sequences are listed in **Fig. S1**). His-tagged versions of the anti-RBD αRep were expressed in *E. coli* and purified. We first explored their affinity for the RBD by biolayer interferometry (BLI) at different concentrations to determine their kinetic rate constants. **Fig. 2** shows the binding of two most potent anti-RBD ligands, αReps C2 and F9. Their affinity for the RBD was about 0.3 and 1.1 nM, respectively. αRep C7 exhibited an affinity in the 10 nM range.

**Fig 2.**
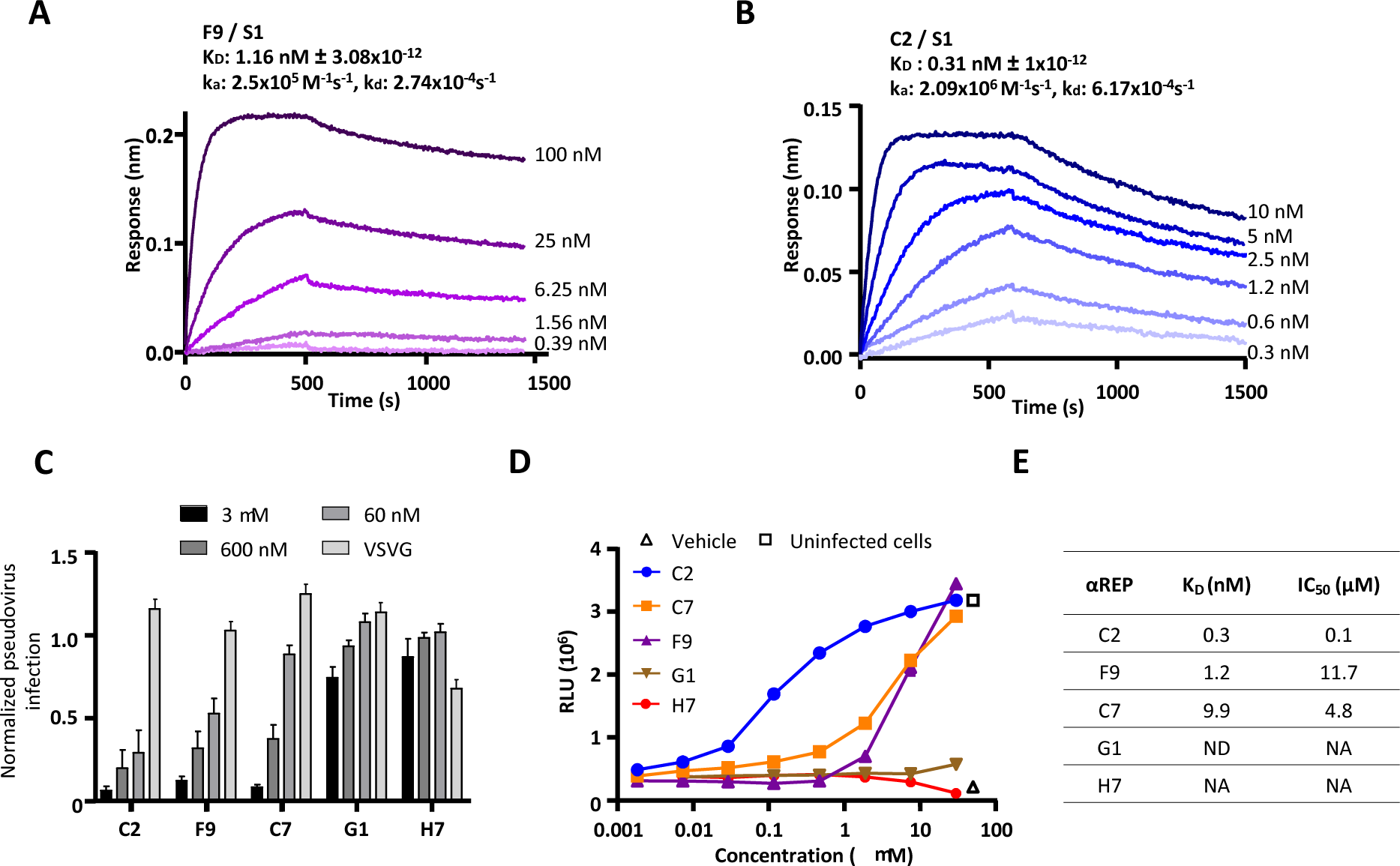
Selection of αReps based on their affinities and neutralization activities. BLI binding kinetics measurements are shown for F9 (**A**) and C2 **(B**). Equilibrium dissociation constants (KD) were determined on the basis of fits, applying 1:1 interaction model; ka, association rate constant; k_d_, dissociation rate constant. **(C)** Pseudo-typed SARS-CoV-2 neutralization assay was shown with selected αRep (C2, F9, C7, G1). An αRep specific to influenza polymerase (H7) was chosen as a negative control. To assess αReps specificity, pseudo-typed VSV-G were incubated with the highest concentration of each αRep (3 µM). Pseudo-type particles entry into cells was quantified by measuring luciferase activity (n=3, mean ± SEM, two-way ANOVA, \**P*<0.05). **(D)** Cell viability of infected cells in presence of dilutions of αReps C2, C7, F9, G1 and H7 was monitored using the CellTiter-Glo Luminescent Assay Kit (Promega). Infected cells (triangle) and mock-infected cells (square) were included in the assay as controls (n=2, mean is presented). **(E)** Half maximal inhibitory concentration (IC_50_) were calculated using “log(inhibitor) vs. normalized response” equation from the neutralization potency curves with GraphPad Prism 8 software. ND: Not done, NA: Not available.

### Identification of neutralizing αReps

We next tested the neutralization activity of the best αREPs candidates against SARS-CoV-2 pseudotyped murine leukemia virions (MLV) as previously described [15]; **Fig. 2C**]. These virions only contain the SARS-CoV-2 spike protein on their surface and behave like their native coronavirus counterparts for entry in cells expressing ACE2. Upon cell entry, the luciferase reporter gets integrated into the host cell genome and is expressed, the measured signal being correlated with αRep neutralization properties. C2, F9 and C7 showed a dose-dependent neutralization activity, C2 displaying the highest neutralisation activity. Neither G1, an additional selected anti-RBD αRep, nor an anti-influenza αREP (H7), used as negative control, displayed notable neutralization activity. None of the αReps tested at the highest concentration (3 µM) displayed neutralization activity against vesicular stomatitis virus G pseudo-typed MLV, demonstrating their specificity.

We confirmed this neutralization activity using SARS-CoV-2 infection of Vero E6 cells (**Fig. 2D**). C2 showed the highest neutralizing potency with a half-maximal inhibitory concentration (IC_50_) value of 0.1 µM, while C7 and F9 αReps displayed IC_50_ values of 4.8 and 11.7 µM, respectively (**Fig. 2E**). G1 as well as the anti-influenza H7 αRep did not show neutralization activity. We thus identified three potent neutralizing αReps, with C2 and F9 displaying affinity in the nM range. These two lasts αReps were retained for further analyses.

### Design of αREP derivates

In order to increase avidity and neutralization activity of these RBD binders, we aimed at generating multivalent αReps. We first determined if F9 and C2 recognized non-overlapping binding sites on the RBD to assess their interest to be linked in a fusion protein. Competitive binding assays carried out by BLI showed that C2 and F9 bindings on the RBD did not interfere in a reciprocal manner (**Fig. 3A and 3B**). Competitive binding assays between these two αReps and soluble hACE2 showed that ACE2 binding occurred efficiently after binding of C2 on the RBD. In contrast, binding of F9 on the RBD partially inhibited recognition of hACE2. As a positive control, VHH72 recognizing the receptor binding motif [16] fully blocked hACE2 binding on the RBD (**Fig. 3C**). These results suggest that the neutralization activity of the C2 αRep is not associated to a steric inhibition of the binding of the RBD on ACE2, and that a fusion between C2 and F9 αReps may be synergistic. We thus engineered bivalent αReps constructs using F9 and C2 αReps. We also generated trivalent αReps through the addition of a trimerization foldon domain (corresponding to the C-terminal part of T4 fibritin) behind C2 and F9 αReps [17].

**Fig 3.**
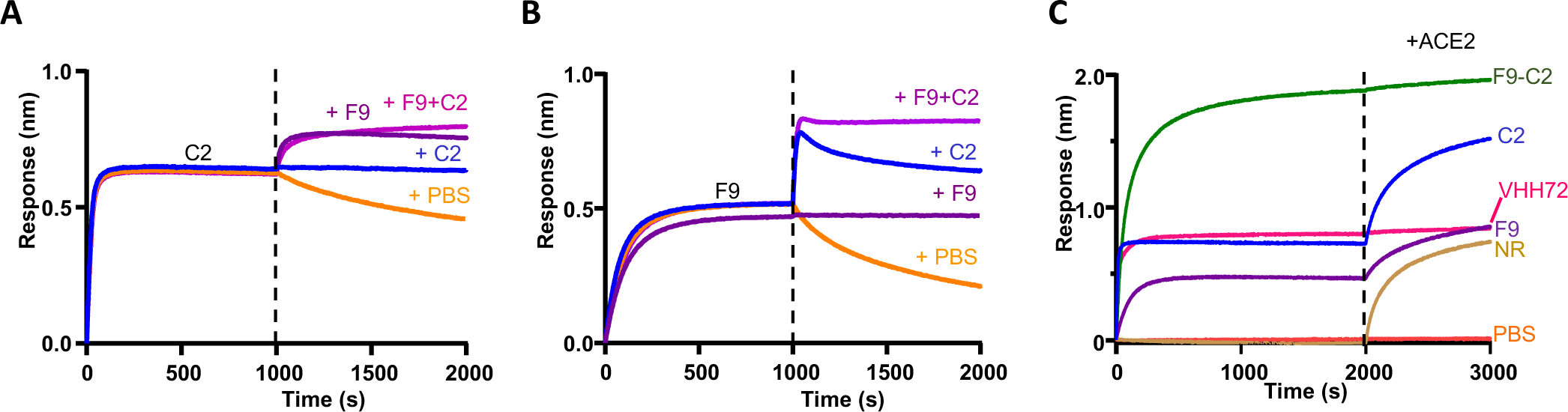
Competitive binding assays. **(A and B)** BLI experiments showed that C2 and F9 could bind RBD simultaneously. **(C)** Binding of ACE2 was assessed after a first association phase with αReps C2 and F9, the F9-C2 construct, the VHH72 [16] or with a negative control (NR). F9-C2 and VHH72 blocked the binding of RBD to ACE2. While F9 inhibited partially ACE2 binding, C2 did not compete with ACE2 binding.

### Properties of αREP heterodimers and homotrimers

To build the F9-C2 and C2-F9 heterodimers, we inserted a 25 amino acid long flexible linker (GGGGS)5 between these two subunits (**Fig. S1**). This linker length (that can reach 8 nm in length) allows the binding of these heterodimers between adjacent RBDs in the trimer, even in the “up” to “down” spike conformers. To generate the homotrimeric C2- and F9-foldon αReps, the foldon sequence was connected to the C-ter of the αREPs through a 16-amino acid long linker (GSAGSAGGSGGAGGSG) (**Fig S1**). These linkers would allow cross-links between spikes at the surface of the virus particle. Unable to express efficiently the C2-F9 construct, only the F9-C2 affinity was characterized by BLI experiments (**Fig. 4A**). F9-C2 displayed an equilibrium dissociation constant (KD) of 91 pM, at least three folds better than that of monomers. F9-C2 also showed a substantially slowed dissociation rate constant of 5.86 x 10^-5^s^-1^ owing to enhanced avidity. Circular dichroism revealed melting temperatures of 86.5°, 88.3° and 86.0°C for C2, F9 and F9-C2, respectively, confirming the high stability of this class of protein (**Fig. S2**).

**Fig 4.**
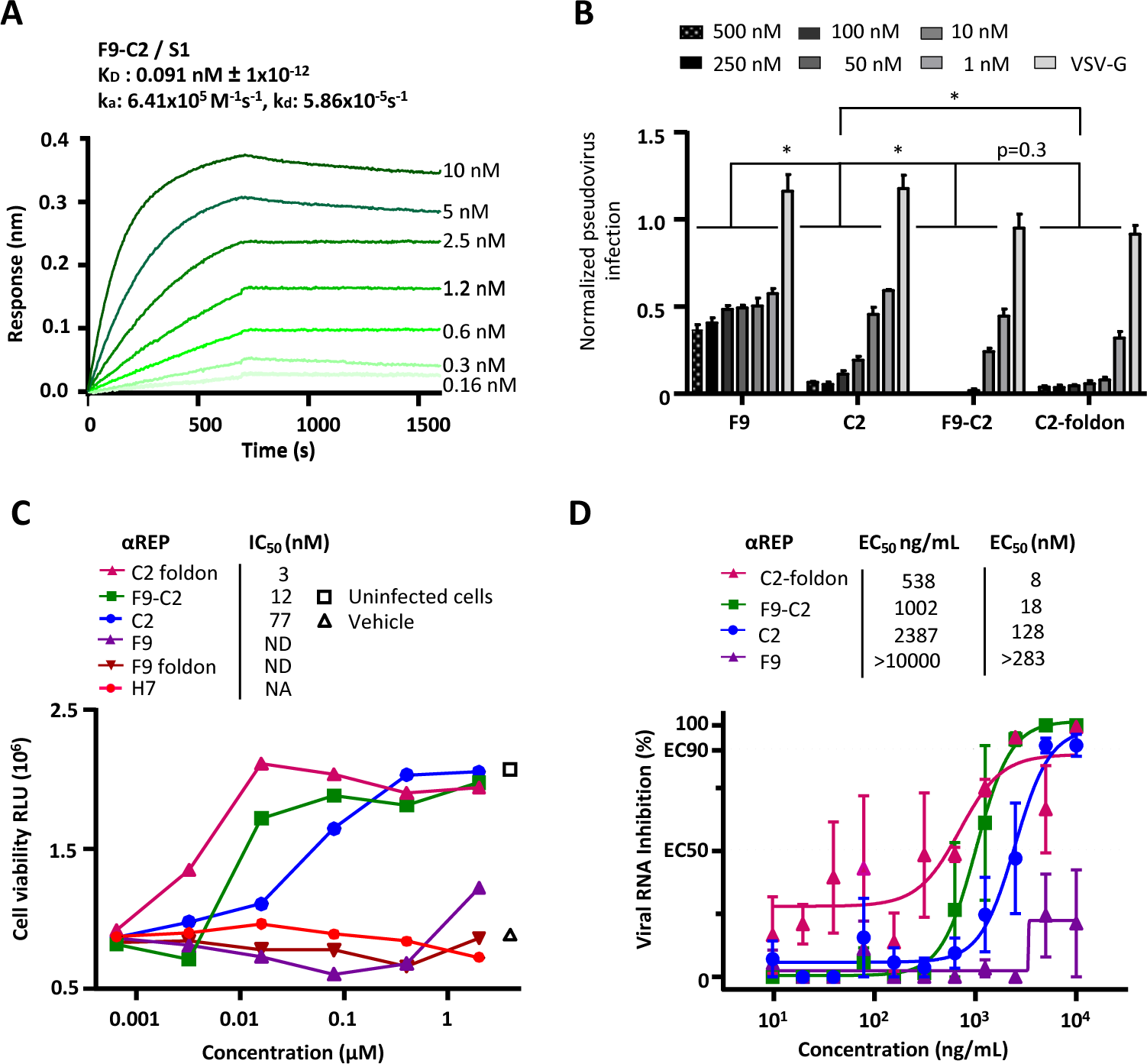
The F9-C2 and C2-foldon constructs properties. **(A)** BLI binding kinetics measurements for F9-C2 to the S1-immobilized biosensor. (**B)** Pseudo-typed SARS-CoV-2 particles neutralization assay was performed with F9, C2, F9-C2 and C2-foldon constructs (n=3, mean ± SEM, two-way ANOVA, \**P*<0.0001). **(C)** Cell viability of SARS-CoV-2-infected cells in presence of dilutions of F9-C2, C2-foldon, C2, F9 and H7 (an αRep negative control) was monitored using the CellTiter-Glo Luminescent Assay Kit (Promega) (n=2, mean is presented). Infected cells (triangle) and mock-infected cells (square) were included in the assay. Half maximal inhibitory concentration (IC_50_) were displayed. **(D)** SARS-CoV-2 neutralisation by aReps constructs. Virus replication was quantified by qRT-PCR in infected cells treated by C2, F9, F9-C2, C2-foldon (n=3, mean ± SEM). Half maximal effective concentration (EC50) were shown.

We next investigated the ability of F9-C2 to block RBD-ACE2 interaction by BLI measurements (**Fig. 3C**). When F9-C2 was bound to the RBD, addition of ACE2 induced no signal shift demonstrating that F9-C2 dimer is a potent inhibitor of spike binding to ACE2, similarly to the VHH72 [16].

We next explored the neutralization activity of F9-C2 and C2- and F9-foldon for comparison with their parental subunits against SARS-CoV-2 spike pseudo-typed MLV (**Fig. 4B**). A synergic effect in neutralisation efficiency was evidenced when the F9 and C2 subunits were covalently linked and when C2 was assembled as a homotrimer. While C2 almost fully blocked entry of SARS-CoV-2 pseudo-type at a concentration of 250 nM, F9-C2 and C2-foldon neutralized infection at 50 nM. We next investigated their viral neutralization potencies in SARS-CoV-2 / Vero E6 cell infection assays by measuring cell viability (**Fig. 4C**) and viral replication (**Fig. 4D**). F9-C2 and C2-foldon were more effective than their monomeric counterparts to protect cells from SARS-CoV-2 infection, with an IC_50_ of 12 nM and 3 nM, respectively, while C2 alone neutralized SARS-CoV-2 with an IC_50_ of 77 nM. F9-foldon displayed a similar activity than its monomeric counterpart indicating no added value of this construction. Quantification of virus replication confirmed the same trend, with an EC_50_ of 18 nM for F9-C2 and 8 nM for C2-foldon, indicating a higher neutralizing activity than C2 (EC50 of 128 nM).

Thus, the covalent linkage between the F9 and C2 subunits or the trimerization of C2 revealed a synergistic effect (∼x 10-25) of αREPs oligomerization to neutralize SARS-CoV-2. Since F9-C2 targeted two different epitopes and may be less sensible to spike antigenic shift, we retained this heterodimer for further *in vivo* analyses.

### F9-C2 prophylaxis limits SARS-CoV-2 infection *in vivo*

In order to evaluate if F9-C2 prophylaxis was effective to limit SARS-CoV-2 infection *in vivo*, we used Syrian golden hamsters known to reflect the infection in human [18]. We focused on the nasal cavity as we choose to examine how a local treatment could limit the start of the infection in a physiological context. We pre-treated the hamsters with 0.6 mg of F9-C2 distributed between the two nostrils 1h prior to infection with 5.10^3^ TCID_50_ of SARS-CoV-2 of the circulating European strain in 2020 (harbouring the D614G mutation in the spike protein) (**Fig. 5A**). After such treatment, we observed the presence of infiltrated αReps on the surface of the epithelium layer, indicating an efficient absorption of the molecule (**Fig. S3**). The group treated with the non-neutralizing αREP G1 lose weight starting from day 2. Treatment with F9-C2 limited weight loss and the difference with G1 treatment almost reach significance at 3 dpi (P=0.057, 2-way ANOVA, **Fig. 5B**). During the 3 days following infection, virus titres in nasal swabs were lower in the group treated with F9 C2 than in the G1-treated group (**Fig. 5C**, two-way ANOVA, *P*<0.0001). In the olfactory turbinates where virus starts to replicate at high titres, amount of viral RNA was significantly lower at 1 dpi in F9-C2 treated animals (Mann Whitney, P=0.0286, **Fig. 5D**), consistent with a tendency of lower expression of inflammation markers, in particular IL-6 and TNFα. At 3 dpi, no significant differences were observed for viral RNA and inflammation markers between the two treatments. Next, we examined if the lower amount of viral RNA at 1 dpi in the nasal cavity of F9-C2-treated animals was reflected at the histological level. While the virus was present in large patches of the epithelium in the G1-treated animals, it was only present in small stretches in F9-C2-treated animals (**Fig. 5E**). We measured the infected area in the rostral part of the nasal cavity at 1 dpi where the infection starts. The difference between G1 and F9-C2 treated animals was close to significance (Mann Whitney, P=0.057, **Fig. 5F**).

**Fig 5.**
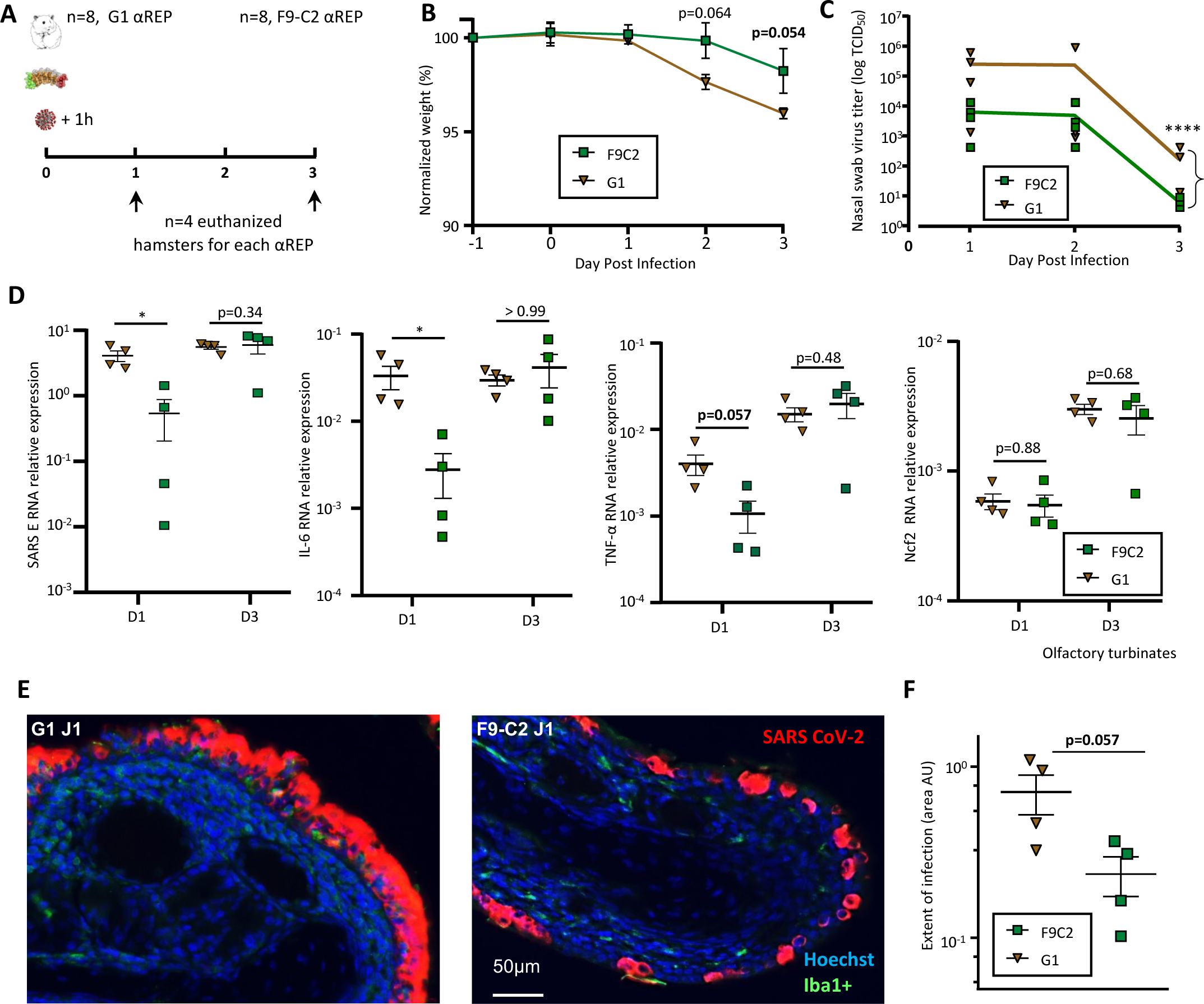
Efficacy of F9-C2 αRep prophylaxis in SARS-CoV-2 infection in a golden Syrian hamster model. **(A)** Overview of the experiment design. 6 mg/kg of αReps were delivered intranasally in hamsters 1h prior to infection with 5.10^3^ TCID_50_ of SARS-CoV-2. (**B**) Evolution of animal weight (n=4, mean of the relative weight to 1-day prior infection ± SEM, two-way ANOVA). (**C**) Evolution of virus titre in nasal swabs (n=4, mean of TCID_50_ ± SEM, two-way ANOVA, \*\*\*\**P*<0.0001) (**D**) Quantification of RNA encoding SARS-CoV-2 protein E, IL-6, TNFα, Ncf2 in the olfactory turbinates, relative to viral infection, inflammation and neutrophil respectively (normalized to ß-actin, mean ± SEM, Mann–Whitney \**P*<0.05). **(E)** Representative images of the infected olfactory epithelium area treated by G1 or F9-C2 in the rostral zone of the nasal cavity (1 dpi) showing respectively a strong and partial infection. **(F)** Measurement of the extent area of infection in the dorso-medial part of the hamster nose. Values represent the mean of infected area (Arbitrary Unit ± SEM, Mann–Whitney \**P*<0.05).

### Repeated F9-C2 treatments further limit SARS-CoV-2 infection *in vivo*

In order to improve the efficiency of the treatment, hamsters were treated with F9-C2 (0.6 mg per dose) 1h prior infection and on days 1 and 2 post-infection (**Fig. 6A**). F9-C2 treatments limited weight loss and the difference reach significance at 3 dpi when compared to controls (P=0.015, 2-way ANOVA, **Fig. 6B**). During the 3 days post-infection, virus titres in nasal swabs were lower in the group treated with F9-C2 than in the control group (**Fig. 6C**, two-way ANOVA, *P*<0.0001). Viral RNA was significantly lower in olfactory turbinates at 1 dpi and 3 dpi in F9-C2 treated animals (Mann Whitney, P=0.0286, **Fig. 6D**) when compared to an irrelevant αRep. This observation correlates with lower expression of inflammation markers (IL-6, TNFα and Ncf2, the last one being related to neutrophil infiltration). We observed less damage of the olfactory epithelium accompanied with a reduction of immune cell infiltration (revealed by the iba1^+^ marker) and desquamated cells in the lumen of the nasal cavity for animals treated by F9-C2, especially at 1 dpi (**Fig. 6E** and **Fig. S4**). The infected area in the rostral part of the nasal cavity at 1 dpi was significantly reduced in F9-C2 treated animals compared to controls (Mann Whitney, P=0.0286, **Fig. 6F**). These results suggested that repeated injections of F9-C2 significantly reduce the spread of the virus up to 3 days post-infection.

**Fig 6.**
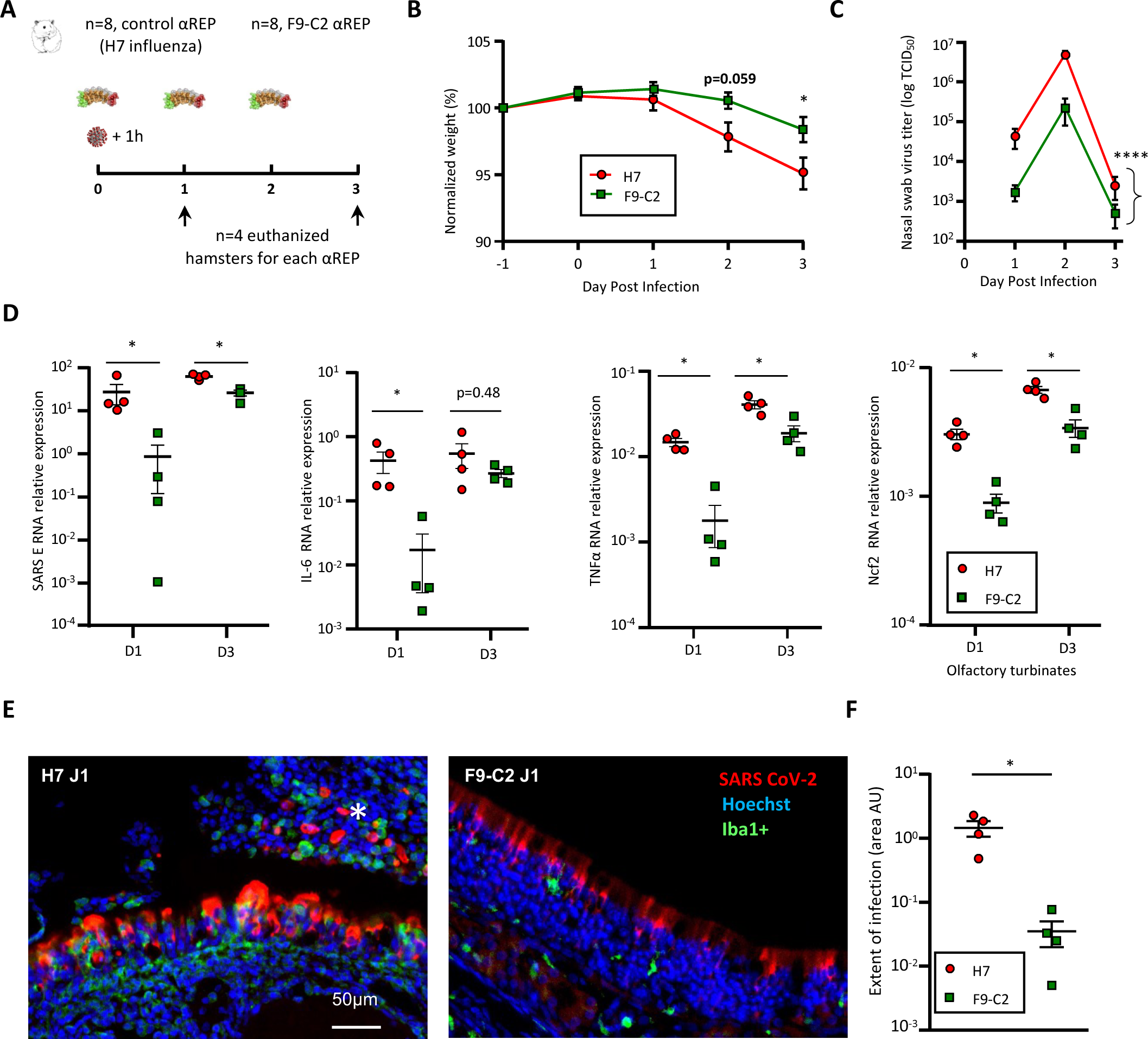
Efficacy of F9-C2 αRep repeated treatments in SARS-CoV-2 infection in a golden Syrian hamster model. **(A)** Overview of the experiment design. 6 mg/kg of αReps were delivered intranasally in hamsters 1h prior to infection with 5.10^3^ TCID_50_ of SARS-CoV-2. The treatment was repeated on 1 dpi and 2 dpi for the group examined at 3 dpi. (**B**) Evolution of animal weight (n=4, mean of the relative weight to 1-day prior infection ± SEM, two-way ANOVA). (**C**) Evolution of virus titre in nasal swabs (n=4, mean of TCID_50_ ± SEM, two-way ANOVA, \*\*\*\**P*<0.0001) (**D**) quantification of RNA encoding SARS-CoV-2 protein E, IL-6, TNFα, Ncf2 in the olfactory turbinates, relative to viral infection, inflammation and neutrophil respectively (normalized to ß-actin, mean ± SEM, Mann–Whitney \**P*<0.05). **(E)** Representative images of the infected olfactory epithelium area treated by F9-C2 or H7 in the dorso-medial zone of the nasal cavity (1 dpi) showing respectively a partial infection with a low number of Iba1^+^ immune cell infiltration and a strong infection associated with damage of the olfactory epithelium and Iba1^+^ cell infiltration as well as desquamated cells in the lumen of the nasal cavity (white asterisk). **(F)** Measurement of the extent area of infection in the dorso-medial part of the hamster nose. Values represent the mean of infected area (Arbitrary Unit ± SEM, Mann–Whitney \**P*<0.05).

### αREP derivates neutralize numerous SARS-CoV-2 variants

We next explored the ability of F9-C2 and C2-foldon to neutralize pseudo-types and SARS-CoV-2 variants. To this end, we first generated five different SARS-CoV-2 S pseudo-typed MLV carrying the RBD mutations specific of each of the α, β, γ, δ and ο variants [19] (**Fig.7A**). In all cases, F9-C2 and C2-foldon appeared to synergise the neutralisation activities of their monomeric counterparts. They inhibited the pseudo-types representatives of the α, β, γ and ο variants entry as efficiently as the parental pseudo-type (with an almost full neutralization of pseudo-typed particles at 100 nM). In contrast, κ / δ like-variant entry was only 90%- and 70%-blocked by F9-C2 and C2-foldon at a concentration of 500 nM. Furthermore, C2-foldon exhibited a similar activity against variants than F9-C2. Figure7B shows the neutralizing potencies of F9-C2 and C2-foldon against authentic virus variants by viral RNA quantification. C2-foldon neutralized efficiently β,γ,δ and ο virus variants (with EC_50_ ranging from 13 to 32 nM)(Fig. 7C). F9-C2 exhibited similar neutralization activities, except for the δ variant (with an EC_50_ >184 nM). As expected, their monomeric counterparts displayed lower specific activities (with EC_50_ > 283 nM).

**Fig 7.**
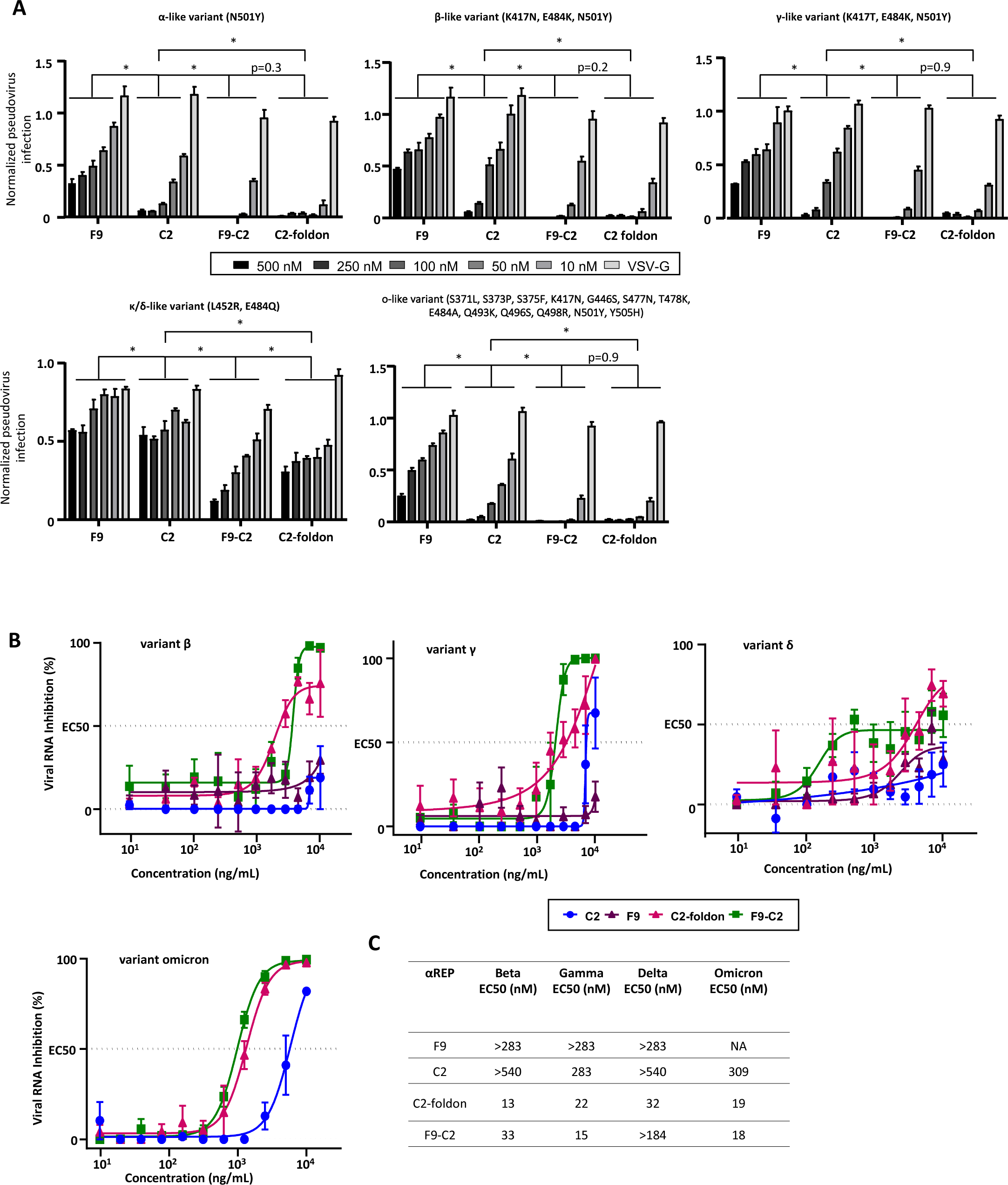
Neutralization activity of the F9-C2 and C2-foldon constructs against SARS-CoV-2 pseudo-typed and virus variants. **(A)** F9, C2, F9-C2 and C2-foldon were tested for their ability to neutralize four SARS-CoV-2 pseudo-typed RBD mutants. Pseudo-typed VSV-G was incubated with the highest concentration of each αRep (500 nM) to validate specificity of αRep neutralization activity (n=3, mean ± SEM, two-way ANOVA, \**P*<0.0001). **(B)** F9, C2, F9-C2 and C2-foldon were tested for their ability to neutralize authentic SARS-CoV-2 virus variants (beta, gamma, delta and omicron) (n=3, mean ± SEM). Chart including the EC_50_ and in nM **(C)** of each αREP for each variant virus is depicted.

## Discussion

From the beginning of the COVID-19 pandemic, numerous studies reported the selection of monoclonal antibodies or nanobodies targeting the SARS-CoV-2 RBD [12] with the aim to block RBD-ACE2 interaction and consequently virus entry in permissive epithelial cells [20], [21]. Although high-affinity antibodies have been prioritized as potential therapeutics, they are expensive to produce in mammalian cell expression systems and they must be injected rather taken orally or by spray [22]. Usually, large doses are required because only a small proportion may cross the epithelial cell barrier lining the airways. Nanobodies represent an interesting alternative to antibodies since they are easy to produce in bacteria or yeast. However, their stability due to structural constraints (the existence of an internal disulfide bridge) in external body compartments may represent a bottleneck for their industrial production and prevent their aerosolization use. In this study, we aimed at producing stable antivirals that could be easily adapted against SARS-CoV-2 variants at low cost.

We used a phage display screening of a library encoding artificial proteins, named αReps, to identify ligands targeting the spike RBD of SARS-CoV-2. Two of them (C2 and F9 αReps) displayed affinity in the nM range. Competitive binding assays showed that these last molecules were able to neutralize the virus through different mechanisms, with C2 binding a site distant of the receptor binding motif to ACE2, in contrast to F9 that compete with ACE2 for binding on RBD. We demonstrated the simplicity of αRep bioengineering to increase the neutralization activity with a multivalent form. The F9-C2 heterodimer and the homotrimeric C2-foldon displayed higher SARS-CoV-2 neutralization activity than the two parental αReps, with IC_50_ of 3 to 12 nM. We explored if nasal instillation of F9-C2 could effectively limit SARS-CoV-2 infection in a hamster model. Such treatment induced a reduction of the virus load in nasal swabs and in the nasal cavity (the primary replication site of SARS-CoV-2), as well as a decline of all the inflammation markers of infection.

Blocking SARS-CoV-2 contamination chains represents a main issue to control Covid-19 pandemic. As the treatment was not sufficient to fully block infection in the nasal cavity, we anticipate that optimisation of the αRep delivery through nebulisation or aerosolization, and using adequate carriers, may increase their efficiency. Indeed, nebulization of a trivalent nanobody improved their effectiveness to reduce the RSV load in nasal swabs in children hospitalized for lower respiratory tract infection [23]. The use of drugs such as αReps derivates or other low-cost antivirals in infected people and contact cases could thus be helpful to block SARS-CoV-2 diffusion in conspecifics.

F9-C2 and C2-foldon resulted in efficient neutralization of a wide variety of SARS-CoV-2 variants (α, β, γ, δ/κ and omicron variants), a feature that may result from the intrinsic high affinity of the αRep subunits for the RBD and their multimerization that allow a less dependence to amino acid substitutions in the target. Current antibody-based therapeutics strategies are being jeopardized by the continuous emergence of SARS-CoV-2 variants through potential loss of binding and neutralization activity [13]. Combining αReps subunits in multivalent forms could thus represent a real option to treat emerging variants.

Beside the fact that αReps assemblies can be easily engineered, i.e. easily associated by linkers or multimerized to generate avidity on targets of interest, these proteins have also favorable biophysical properties of production, purification and stability and can be very efficiently produced (20 mg of purified protein C2, F9, F9-C2 and C2-foldon per liter of bacterial culture) by recombinant protein expression technologies in bacteria. They are particularly robust, highly thermostable and can be stored at room temperature, which is a significant advantage for further therapeutic developments.

Immunogenicity of the αReps is a potential problem that should be addressed in the future, but their relatively small size (the C2 αRep is 18.5 kDa), their association through flexible linkers and their high solubility and stability may result in low immunogenic activity and may not induce adverse undesirable effects when delivered in the nasal cavity.

To conclude, we selected artificial proteins (αReps) as specific and versatile neutralizing binders targeting the spike of SARS-CoV-2. These biosynthetic proteins provide starting points for SARS-CoV-2 therapeutics able to target emerging variants. With technical optimisation in binder selection and effort to stabilize them in the nasal cavity, we believe that stable proteinaceous inhibitors like αReps and derivates have a real future to threat future pandemics associated to various emerging respiratory viruses.

## Materials and methods

### Production of the receptor binding domain (RBD) of the SARS-CoV-2 spike

The RBD (223 amino acids starting at position 319 of the spike sequence) coding sequence was cloned in frame behind a sequence encoding a signal peptide and in front of a His-tag coding sequence in the eukaryotic pYD11 expression plasmid. The resulting plasmid was transfected with PEImax (24765-1) (Polysciences, Inc.) in EXPI-293F cells (A14527) (Thermofisher). Transfected cells were then maintained in EXPI expression medium (Gibco, Thermofisher). Cells were removed by mild centrifugation at day+7, and the RBD was extracted from the cell culture medium by Ni^2+^ affinity chromatography followed by gel filtration. About 10 mg of RBD were purified for αReps screening and further characterization.

### Screening of the αRep library against the RBD

The construction of the αRep phage library 2.1 has been previously described [24]. The αRep library was constructed by polymerization of synthetic microgenes corresponding to individual HEAT-like repeats, and the αRep proteins were expressed at the surface of M13-derived filamentous phages. The library is estimated to contain 1.7 x 10^9^ independent clones. The αRep library screening was carried out as described by Hadpech et al., 2017 [8] with minor modifications. Purified RBD diluted at 1 mM in PBS containing 0.05% Tween-20 (PBST) was immobilized on Ni^++^-NTA-biotin streptavidin-coated 96-well ELISA plate by incubation overnight at 4°C in a moisture chamber. The coated wells were washed four times with PBST, and saturated with blocking solution (2% BSA in PBST; 200 µL/well) for 1h, after which an aliquot of the phage library was added to the RBD coated wells, and incubated at room temperature for 2h with shaking. Next, wells were extensively washed in PBST, and bound phages were eluted by three successive rounds of adsorption/elution. Phage elution was performed by an acidic glycine solution (0.1 M glycine-HCl, pH 2.5) and buffered using Tris 1 M. The population of αRep-displayed phages eluted from the RBD bait was amplified and subcloned in a XL-1 Blue cells. Individual phage clones were selected and amplified as previously described [5], and their respective binding activity towards the RBD was determined by ELISA. 100 μL-aliquots of purified RBD were diluted in PBS and loaded into the wells of a Ni^++^-NTA-biotin streptavidin-coated ELISA plate, then incubated overnight at 4°C. The coated plate was washed four times with TBST, then blocked with PBST-BSA (200 μL per well) for 1 hour with shaking. After a washing step, 100 μL-aliquots of each phage culture supernatant were added to the wells and incubated at room temperature for 1 hour, followed by HRP-conjugated mouse anti-M13 (Interchim) diluted to 1:2,000 in TBST-BSA (100 μL-aliquot per well), and incubation proceeded for an extra 1 hour. The wells were washed again, prior to the addition of 100 μL BM Blue POD soluble Substrate (Roche). Reaction was stopped with 1 N HCl, and absorbance measured at 450 nm. Phage clones showing a high binding activity towards the immobilized target were sequenced and kept for cytoplasmic expression of individual αRep proteins.

### αReps expression and purification

The αRep genes encoding the RBD binders were subcloned in the bacterial expression vector pQE81 and resulting plasmids used for transforming Rosetta cells. αRep gene expression was induced by IPTG addition (0.5 to 1 mM final concentration) for 4 hours at 37°C. Next, bacteria were pelleted by centrifugation (5.000 x g for 30 min at 4°C) and bacterial cell pellets were resuspended in 200 mM NaCl, 20 mM Tris pH 7.4 to 8, containing a cocktail of protease inhibitors (Roche Diagnostics GmbH). Then, bacterial suspension was lysed by sonication in ice (5 times x (30 s sonication at around 40% sonication amplitude and 30 s rest)) using a Q700 QSONICA Sonicator. Bacterial cells lysates were clarified by centrifugation at 10.000 × g for 30 min at 4°C. Soluble αReps were purified by affinity chromatography on HisTrap columns (GE Healthcare Life Sciences) and analyzed by SDS-PAGE. Fractions of interest were pooled and injected in a Gel filtration Superdex S200 previously equilibrated with PBS. Fractions containing purified αReps were pooled and frozen at -20°C.

### αReps circular dichroism

Circular dichroïsm spectra were recorded with a Jasco J-810 system. 200 μL of each purified αReps C2, F9 and F9-C2 in PBS buffer at respectively 0.5 mg/mL, 0.75 mg/mL and 1 mg/mL were disposed in a 3 mm quartz cuvette. Samples were exposed to increasing temperatures from 25°C to 95°C with a measurement every 0.5°C at 230 nm.

### Affinity determination by Bio Layer Interferometry

Binding kinetics experiments were performed on an Octet system (Octet RED96) (FortéBio, CA). A black bottom 96-well microplate (Greiner Bio-One # 655209) was filled with 200 μL of solution (αReps in PBS buffer) and agitated at 1 000 rpm, and all experiments were carried out at 25°C. Tips were hydrated in PBS buffer for 1 hour at room temperature prior experiments. Biotinylated SARS-CoV-2 RBD or S1 (4 µg/mL) were loaded on streptavidin SA (18-0009) biosensors (Pall ForteBio) for 1 min. After a baseline step in assay buffer (PBS [pH 7.4], 0.1% bovine serum albumin, 0.02% Tween 20) and a quenching step in 5 μg/mL biocytin, ligand-loaded sensors were dipped into known concentrations of αReps for an association phase during 500 to 700 seconds. The sensors were then dipped in assay buffer for a dissociation step during 1000 seconds in assay buffer. Association and dissociation curves were globally fitted to a 1:1 binding model except for F9-C2 whose fitting model was 1:2. Binding curves were fit using the “association then dissociation” equation in the FortéBio Data analysis software version 7.1 to calculate K_D_.

### Competition assays

Competition assays between αReps were performed with biotinylated SARS-CoV-2 RBD (4 μg/mL) loaded on SA biosensors for 1 min. After a baseline step in assay buffer and a quenching step, a first association is realised with an αRep at 100 nM for 2000 seconds followed by a second association step with a second αRep at 100 nM during 1000 seconds, and finally a short dissociation step of 300 sin assay buffer. For competition between αREP and ACE2, the second analyte is replaced by soluble ACE2 at 100 nM.

### MLV pseudo-typed particles production and αReps neutralization activity of SARS-CoV-2 pseudotypes

For pseudotyping, murine leukemia virus pseudo-typed particles (PP) containing the spike of SARS-CoV-2 or derived mutants were produced according to a published protocol [15]. Briefly, HEK-293TT cells (10^6^ cells per P6 well) were transfected with plasmids encoding GAG-POL, F-LUC and SARS-CoV-2 spikes. Supernatants containing the pseudo-typed particles were harvested at 48 hours after transfection, pooled and filtered through 0.45 μm pore-sized membranes. Five different PP were produced, a first containing the spike of the SARS-CoV-2 type (Genebank accession number: MN908947), and the four others containing the spike with RBD mutations representatives of the α variant (N501Y substitution), the β variant (N501Y, K417N and E484K mutations), the γ variant (N501Y, K417T and E484K substitutions), the κ/δ variant (substitutions L452R and E484Q) and the ο variant (S371L, S373P, S375F, K417N, N440K, G446S, S477N, T478K, E484A, Q493K, G496S, Q498R, N501Y, Y505H substitutions).

The day before transduction, around 20.000 hACE2-HEK-293T cells were seeded in wells of P96 plates. Three to five-folds serial dilutions of αReps in complete medium (DMEM + 10% FBS) were pre-incubated with pseudo-typed particles to a final volume of 200 μL for one hour at 37°C. The mixture was next added to cells for 48 to 72 hours at 37°C. Then, cells were washed in PBS and lysed as indicated by the manufacturer (Promega # E1501 Luciferase Assay system). Luciferase activity was measured on an Infinite M200Pro TECAN apparatus. MLV pseudo-typed particles with the G glycoprotein of the vesicular stomatitis virus (VSV) were used to monitor the specificity of the neutralization activity at the highest αRep concentration used in the assay. Luciferase activity of each condition was normalised to the reference value, i.e the luciferase activity of the infected cells without αReps.

### Virus stock production and quantification of αReps neutralization activity

SARS-CoV-2 isolate France/IDF0372/2020 was kindly supplied by Sylvie van der Werf. Viral stocks were prepared by propagation in Vero E6 cells in Dulbecco’s modified Eagle’s medium (DMEM) supplemented with 2% (v/v) fetal bovine serum (FBS, Invitrogen). Titres of virus stocks were determined by plaque assay.

To measure the neutralization activity of αReps, three or five-folds dilutions of αReps were mixed with 2 x 10^4^ plaque forming units of SARS-CoV-2 in DMEM for one hour at 37°C. This mixture was added to Vero-E6 cells (CRL-1586, ATCC) seeded in a 96-well plate one day before. Cell viability was measured 72 hours post-infection by adding 100μl of CellTiter-Glo reagent to each well as described in the manufacturers protocol (CellTiter-Glo Luminescent Cell Viability Assay, Promega # G7571). Luminescence was quantified using the Infinite M200Pro TECAN apparatus.

### Golden Syrian hamster infections and assessment of αREPs antiviral activity

#### Hamster infections

Thirty-two specific-pathogen-free (SPF) 8 weeks-old Golden Syrian hamsters (*Mesocricetus auratus*, males, provided by Janvier-Labs, Le Genest-Saint-Isle, France) housed under BSL-III conditions were kept according to the standards of French law for animal experimentation. The study was carried out following a protocol approved by the ANSES/EnvA/UPEC Ethics Committee (CE2A-16) and authorized by the French ministry of Research under the number APAFIS#25384-2020041515287655 v6 in accordance with the French and European regulations.

Groups of eight hamsters were treated with either F9-C2, G1 or H7 (recognizing an influenza virus protein) by nasal instillation (80 µL of PBS containing 0.6 mg of αRep). Animals were infected 1 hour later (**Fig. 5A** and **6A**) with 5 .10^3^ TCID_50_ of SARS-CoV-2 isolate BetaCoV/France/IDF/200107/2020 kindly provided by the Urgent Response to Biological Threats (CIBU) hosted by Institut Pasteur (Paris, France) and headed by Dr. Jean-Claude Manuguerra. The human sample from which strain BetaCoV/France/IDF/200107/2020 was isolated has been provided by Dr Olivier Paccoud from the La Pitié-Salpétrière Hospital. With a history of 3 passages in Vero E6 cells, the seed stock was titrated in Vero E6 cells to a concentration of 6.8 log10 TCID_50_/mL. Nasal swabs were performed daily to measure the secreted virus load by brushing the nostrils of the animal. Eight animals were euthanized 1-day post infection (dpi). The treatment with αReps was repeated daily for one group of hamsters euthanized at 3 dpi. For each hamster, we collected the head which was separated into two hemi-heads, of which one was used for histology and the other for qPCR analysis.

#### Gene expression quantification by RT-qPCR

Total RNA was extracted from frozen olfactory mucosa and lungs using the Trizol method. Oligo-dT first strand cDNA were synthesized from 5 μg total RNA by iScriptAdv cDNA kit (Biorad, # 1725038) following the manufacturers recommendations. qPCR was performed using cDNA templates (5 μL) added to a 15 μL reaction mixture containing 500 nM primers (sequences in Table 1) and iTaq Universal SYBR Green Supermix (Biorad, # 1725124) using a thermocycler (Mastercycler ep realplex^2^, Eppendorf). The expression levels of target genes were measured using the Eppendorf realplex^2^ software. A dissociation curve was carried out at the end of the PCR cycle to verify the efficiency of the primers to produce a single and specific PCR amplification. Quantification was achieved using the ΔΔCt method. Standard controls of specificity and efficiency of the qPCR assays were performed. The mRNA expression was normalized to the expression level of β-actin and an efficiency corrective factor was applied for each primer pair [25].

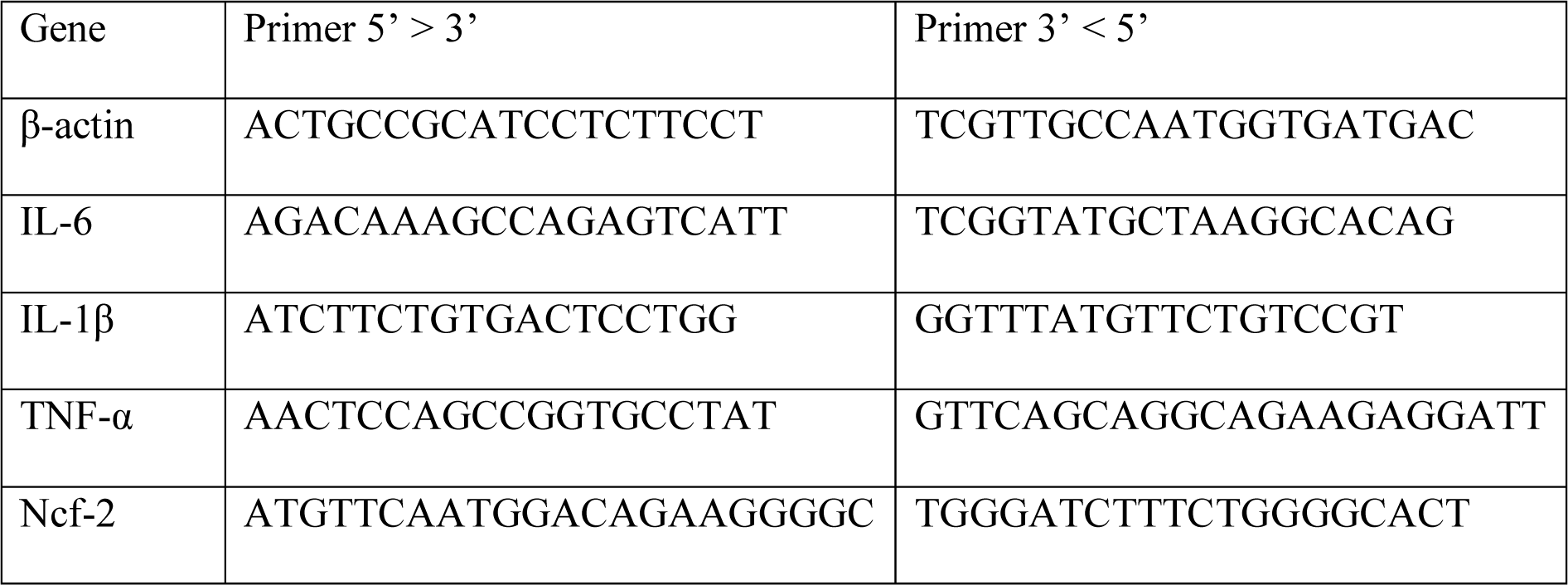
Primers used for qPCR reactions.

#### Viral titration of nasal swabs

Naal swabs were diluted in 400 μL of DMEM medium supplemented with 1% sodium pyruvate and antibiotics and stored at -80°C until titration by tissue culture infectious dose 50% (TCID_50_) on Vero E6 cells. Briefly, each nasal swab was incubated in eight consecutive wells of 96-well microplates, and then serially diluted from 10^-1^ to 10^-6^ within DMEM containing 1% Sodium Pyruvate and 1% antibiotics (Penicillin/Streptomycin). After 1h30 incubation at 37°C, 100μL of complete DMEM medium with 5% FCS, 1% sodium pyruvate and antibiotics are added. The cells are then incubated for 4 days at 37°C. Then, microplates were qualitatively read according to an “all or nothing” scoring method for the presence of viral cytopathic effect (CPE). Infectious titres are expressed as TCID_50_ per mL according to the Spearman Karber method [26].

#### Histopathology

The immunohistochemistry analysis of the olfactory mucosa tissue sections was performed as described previously in mice [27]. Briefly, the hamster hemi-head was fixed for 3 days at room temperature in 4% Neutral Buffer Formalin (F0043, Diapath), then decalcified for three weeks in Osteosoft mild decalcifier solution (1017281000, Merck) at 4°C. Blocks and tissues were cryoprotected with sucrose (30%) then embedding with Epredia Neg-50 (11912365, FisherScientific). Cryo-sectioning (14 μm) was performed on median transversal sections of the nasal cavity, perpendicular to the hard palate, in order to highlight vomeronasal organ, olfactory mucosa, Steno’s gland and olfactory bulb. Sections were stored at −80°C until use. Sections were rehydrated in PBS for 5 min and non-specific staining was blocked by incubation in PBS with 1% bovine serum albumin (BSA) and 0.05% Tween-20. The sections were then incubated overnight in PBS with 0.2% BSA and 0.05% Tween-20 with primary antibodies directed against SARS-CoV-2 Nucleocapsid Protein (1:500; mouse monoclonal, # ZMS1075, Merck); Iba1 (1: 1000; goat polyclonal; ab178846, Abcam, France). Fluorescence staining was performed using goat anti-rabbit Alexa-Fluor-488 (1:800; Molecular Probes – A32731; Invitrogen, Cergy Pontoise, France) and donkey anti-mouse Alexa-Fluor 555 (1:800; Molecular Probes – A32773; Invitrogen, Cergy Pontoise, France) secondary antibodies. Images were taken at ×100 magnification using an Olympus IX71 inverted microscope equipped with an Orca ER Hamamatsu cooled CCD camera (Hamamatsu Photonics France, Massy, France). Images were quantified using ImageJ (Rasband, W.S., ImageJ, U. S. National Institutes of Health, Bethesda, Maryland, USA, http://imagej.nih.gov/ij/, 1997–2012) to threshold specific staining of SARS-CoV-2 in the dorso-medial area of the nasal cavity. We measured the total infected area in this zone displaying the highest area of olfactory epithelium using the same threshold for all animals.

### Quantification of αReps neutralization activity of SARS-CoV-2 virus variants

#### Cell line

VeroE6 TMPRSS2 cells (ID 100978) were obtained from CFAR and were grown in minimal essential medium (Life Technologies) with 7 .5% heat-inactivated fetal calf serum (FCS; Life Technologies with 1% penicillin/streptomycin (PS, 5000U.mL−1 and 5000µg.mL−1 respectively; Life Technologies) and supplemented with 1 % non-essential amino acids (Life Technologies) and G-418 (Life Technologies), at 37°C with 5% CO2.

#### Virus strains

SARS-CoV-2 strain BavPat1 was obtained from Pr. C. Drosten through EVA GLOBAL (https://www.european-virus-archive.com/) and contains the D614G mutation. SARS-CoV-2 Alpha, (201/501YV.1) was isolated from a 18 years-old patient. The full genome sequence has been deposited on GISAID: EPI_ISL_918165. The strain is available through EVA GLOBAL: UVE/SARS-CoV-2/2021/FR/7b (lineage B 1. 1 .7, ex UK) at https://www.european-virus-archive.com/virus/sars-cov-2-uvesars-cov-22021fr7b-lineage-b-1-1-7-ex-uk. SARS CoV-2 Beta (SA lineage B 1.351) was isolated in France in 2021, The strain is available through EVA GLOBAL: UVE/SARS-CoV-2/2021/FR/1299-ex SA (lineage B 1.351) at https://www.european-virus-archive.com/virus/sars-cov-2-uvesars-cov-22021fr1299-ex-sa-lineage-b-1351. Sars-Cov-2 Gamma (SARS-CoV-2/2021/JP/TY7-503 lineage P.1, ex Brazil) was isolated in Japan in January 2021. The full genome sequence has been deposited on GISAID: EPI_ISL_877769. The strain is available through EVA GLOBAL at https://www.european-virus-archive.com/virus/sars-cov-2-virus-strain-sars-cov-22021jpty7-503-lineage-p1-ex-brazil. SARS-CoV-2 Delta, (India lineage B.1.617.2): the full genome sequence has been deposited on GISAID: EPI_ISL_2838050. The strain is available through EVA GLOBAL: SARS-CoV-2/2021/FR/0610 (Lineage B 1.617.2, Delta) at https://www.european-virus-archive.com/virus/sars-cov-2-virus-strain-sars-cov-22021fr0610-lineage-b-16172-delta.

To prepare the virus working stocks, a 25cm2 culture flask of confluent VeroE6 TMPRSS2 cells growing with MEM medium with 2.5% FCS was inoculated at a multiplicity of infection (MOI) of 0.001. Cell supernatant medium was harvested at the peak of replication and supplemented with 25 mM HEPES (Sigma-Aldrich) before being stored frozen in aliquots at - 80°C. All experiments with infectious virus were conducted in a biosafety level 3 laboratory.

#### EC50 and EC90 determination

One day prior to infection, 5×10^4^ VeroE6/TMPRSS2 cells per well were seeded in 100 µL assay medium (containing 2.5% FCS) in 96 well culture plates. αReps were diluted in PBS with ½ dilutions from 10.000 to 9.76 ng/ml. The next day, 25 µL of a virus mix diluted in medium was added to the wells. The amount of virus working stock used was calibrated prior to the assay, based on a replication kinetics, so that the viral replication was still in the exponential growth phase for the readout as previously described [28–32]. This corresponds here to a MOI of 0.002. Then eleven 2-fold serial dilutions of αReps in triplicate were added to the cells (25 µL/well, in assay medium). Four virus control wells were supplemented with 25 µL of assay medium. Plates were first incubated 15 min at room temperature and then 2 days at 37°C prior to quantification of the viral genome by real-time RT-PCR. To do so, 100 µL of viral supernatant was collected in S-Block (Qiagen) previously loaded with VXL lysis buffer containing proteinase K and RNA carrier. RNA extraction was performed using the Qiacube HT automat and the QIAamp 96 DNA kit HT following manufacturer instructions. Viral RNA was quantified by real-time RT-qPCR (GoTaq 1-step qRt-PCR, Promega) using 3.8 µL of extracted RNA and 6.2 µL of RT-qPCR mix and standard fast cycling parameters, i.e., 10 min at 50°C, 2 min at 95°C, and 40 amplification cycles (95°C for 3 sec followed by 30 sec at 60°C). Quantification was provided by four 2 log serial dilutions of an appropriate T7-generated synthetic RNA standard of known quantities (102 to 108 copies/reaction). RT-qPCR reactions were performed on QuantStudio 12K Flex Real-Time PCR System (Applied Biosystems) and analyzed using QuantStudio 12K Flex Applied Biosystems software v1.2.3. Primers and probe sequences, which target SARS-CoV-2 N gene, were: Fw: GGCCGCAAATTGCACAAT; Rev : CCAATGCGCGACATTCC; Probe: FAM-CCCCCAGCGCTTCAGCGTTCT-BHQ1. Viral inhibition was calculated as follow: 100* (quantity mean VC-sample quantity)/ quantity mean VC . The 50% and 90% effective concentrations (EC50,; compound concentration required to inhibit viral RNA replication by 50%) were determined using logarithmic interpolation after performing a nonlinear regression (log(agonist) vs. response -- Variable slope (four parameters)) as previously described [28–32]. All data obtained were analyzed using GraphPad Prism 8 software (Graphpad software).

### Statistical analysis

Data shown as the means ± SEM. All statistical comparisons were performed using Prism 8 (GraphPad). Quantitative data were compared across groups using two-way ANOVA test for pseudovirus assay, weight, nasal swab and virus titre evolution. All other parameters were tested using the Mann-Whitney non-parametric test. Statistical significance was determined as p-value <0.05.

## Acknowledgments

We thank Sylvie van der Werf for the gift of the SARS-CoV-2 strain France/IDF0372/2020, and Ameline Batsché for her help in αRep production and purification.

## Author contributions

S.T.: Methodology, Formal analysis, Investigation, Data curation, Writing – original draft, Writing – review & editing, Visualization, and Supervision. N.L.: Investigation and Data curation. A.D.: Investigation and Data curation. A.U.: Resources, Methodology, Investigation, Writing – review & editing. M.V.-L.: Resources, Methodology, Investigation, Writing – review & editing. M.L.: Methodology, Data curation, Writing – review & editing. B.d.C.: Investigation and Data curation. C.B.: Investigation and Data curation. A.D.: Investigation and Data curation. A.F.: Investigation and Data curation. A.S.-A.-D.: Investigation and Data curation. J. M.: Methodology, Investigation and Data curation. B.K.: Resources. J.D.: Formal analysis and Data curation. A.R.: Methodology, Formal analysis, Investigation, Data curation, Writing – review & editing, Visualization, and Supervision. P.M.: Methodology, Formal analysis, Investigation, Data curation, Writing – review & editing, Visualization, and Supervision. S.L.P.: Methodology, Formal analysis, Investigation, Data curation, Writing – review & editing, Visualization, and Supervision. N.M.: Methodology, Formal analysis, Investigation, Data curation, Writing – original draft, Writing – review & editing, Visualization, and Supervision. B.D.: Conceptualization, Methodology, Formal analysis, Investigation, Data curation, Writing – original draft; Writing – review & editing, Supervision, and Funding acquisition.

## Data and materials availability

αRep sequences are shown in **Fig S1**.

## Supplementary Figures

**Supplementary Fig. S1.**
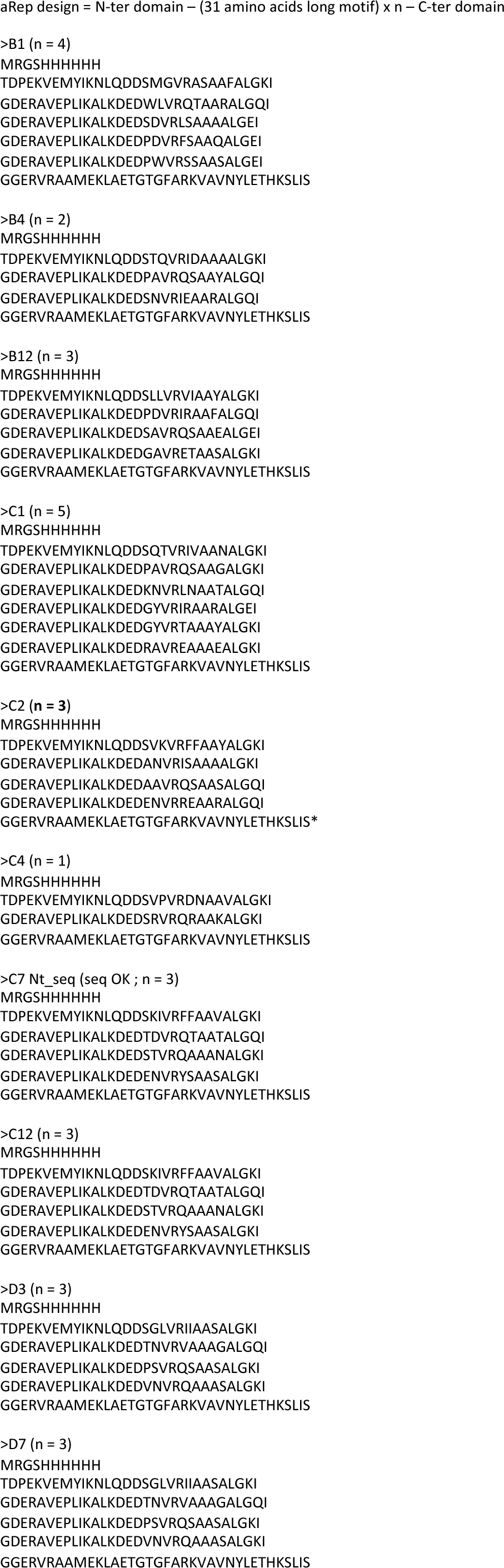
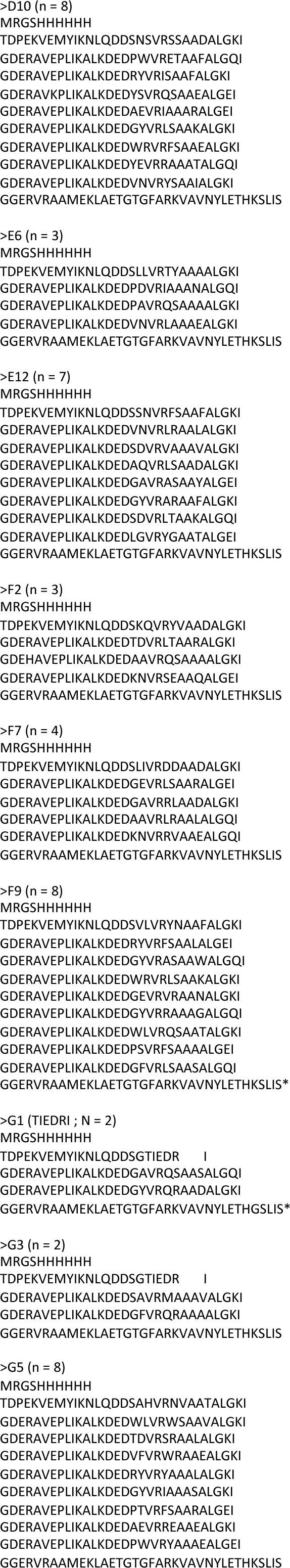
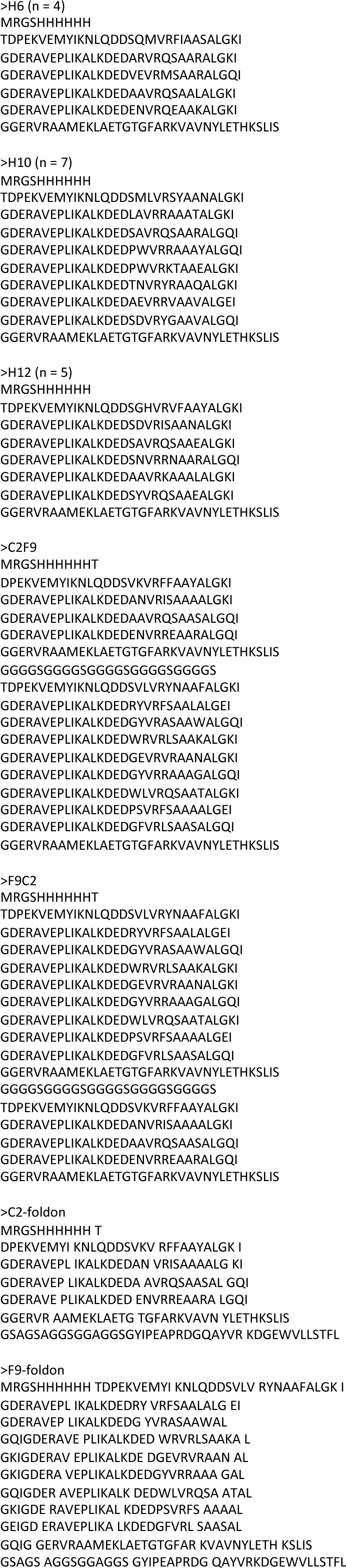
αReps sequences

**Supplementary Fig. S2.**
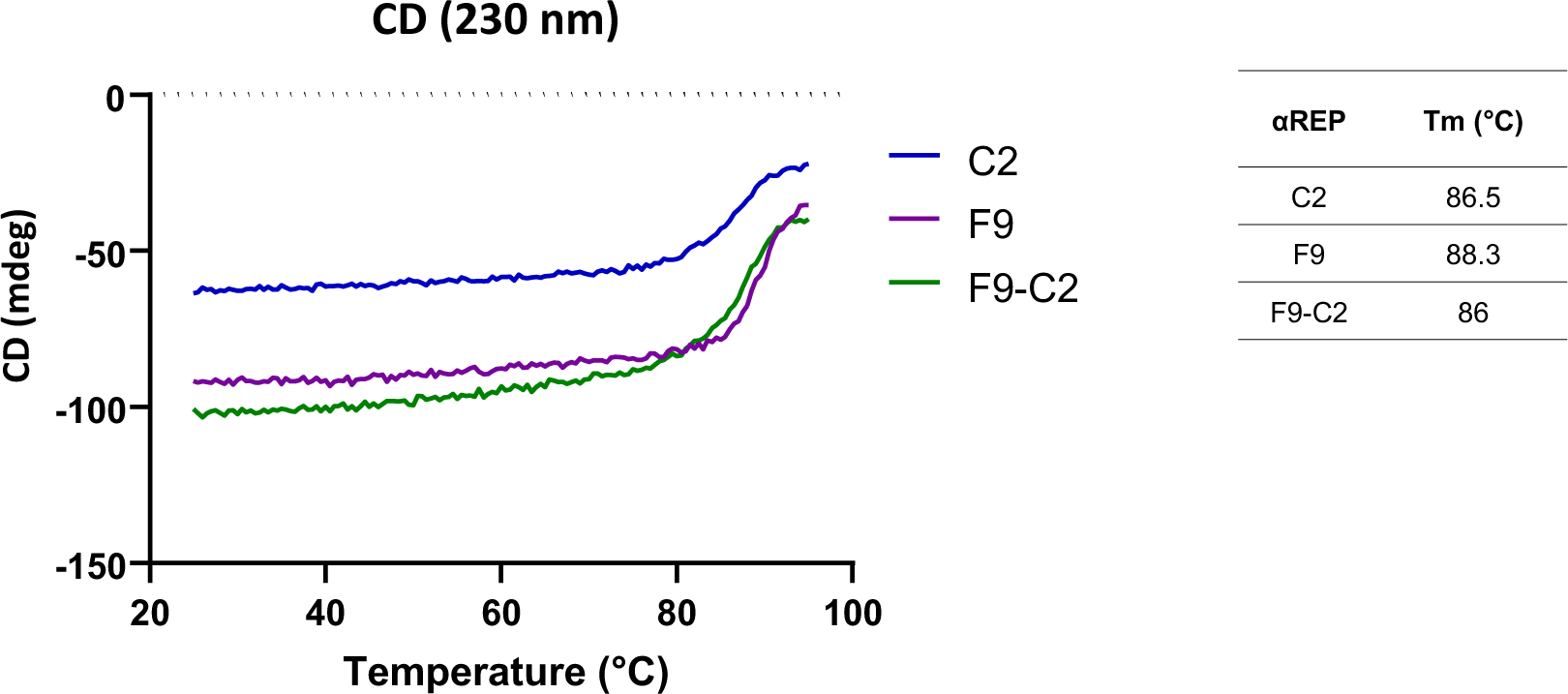
Thermal denaturation of αReps assessed by circular dichroism measurement of molar ellipticity at 230 nm.

**Supplementary Fig. S3.**
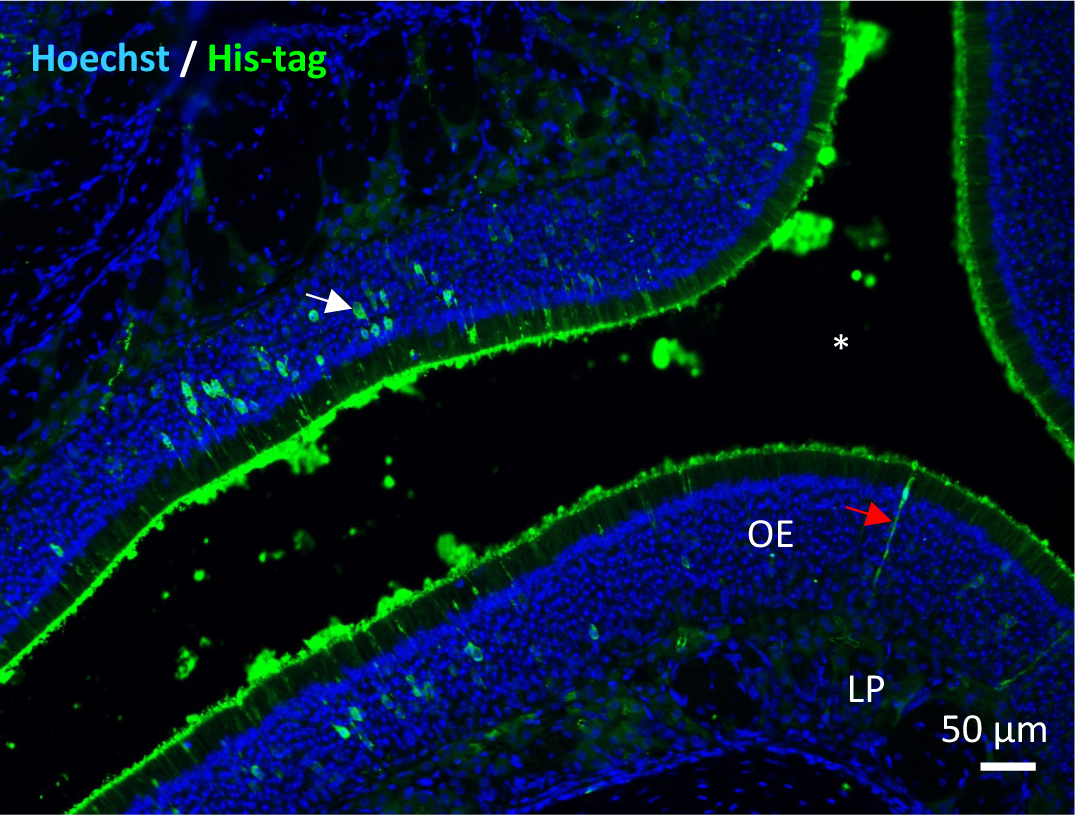
Representative image of the olfactory epithelium (dorsomedial area) 15 min after F9-C2 instillation revealed by an anti-His Tag. F9-C2 is mainly present in the mucus layer but some cells have integrated them (OE: Olfactory Epithelium / LP: Lamina Propria / white asterisk Lumen of the nasal cavity / red arrow: sustentacular cell like shape / white arrow (olfactory sensory neuron like shape).

**Supplementary Fig. S4.**
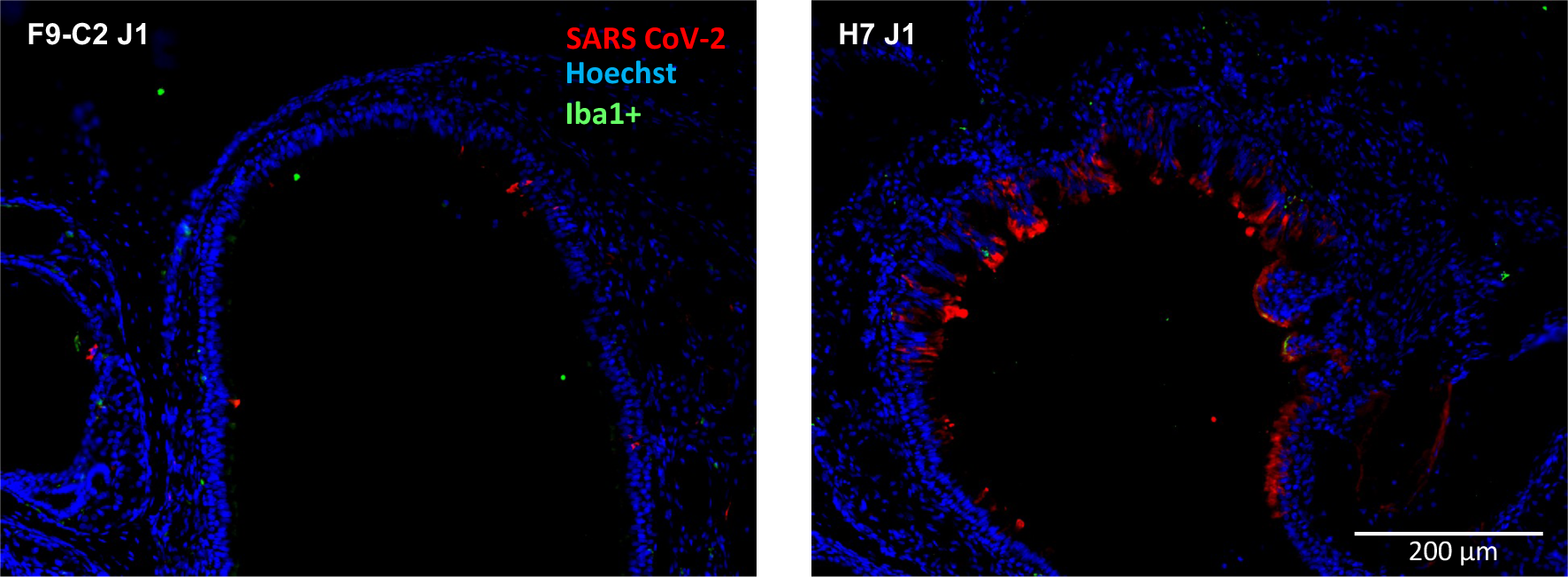
Representative images of the dorso-medial part of the infected hamster nose treated by F9-C2 or H7 (1 dpi). SARS-CoV-2 infected cells were revealed with an anti-N antibody. F9-C2 was found to protect the nasal cavity epithelium.

